# Structural Basis of Asymmetric Class C GPCR activation and distinct Gq coupling induced by force stimulation

**DOI:** 10.64898/2025.12.08.692793

**Authors:** Jin-Hui Ding, Jia-Le Wang, Wei-Feng Zhang, Shu-Hua Zhou, Chan-Juan Xu, Wei Ding, Wan-Ping Zhuang, Qi-Yue Zhang, Chao Zhang, Jie Cheng, Yu-Nan Sun, Yu Sun, Ren-Jie Chai, Xiao Yu, Zhao Yang, Jian-Feng Liu, Jin-Peng Sun

## Abstract

Mechanosensation is essential for diverse physiological processes. While many G protein-coupled receptors (GPCRs) are known to be mechanosensitive, the underlying mechanism remains largely unknown. Here, we reveal that mechanical force induces Gq activation of mGlu2, a balance modulator residing in kinocilia, but not other 7 mGlu members. Force activates mGlu2 through a conserved N-terminal force transduction motif (FTM) via a unique *cis* mechanism. We engineered a potent FTM-derived peptide agonist that recapitulates force-induced activation and resolved cryo-EM structures of apo-mGlu2, FTM-mGlu2, LFTM-ΨEK-mGlu2 and LFTM-ΨEK-mGlu2-Gq. The structures reveal that an atypical FTM binding to a previously uncharacterized pocket induces asymmetric 7TM domain rearrangement, enabling Gq coupling via an ICL1-TM3/6/7 interface, fundamentally distinct from the glutamate-induced Gi coupling mode of mGlu2. Compared with Gi-coupled mGlu2, the α5 helix of Gq rotated by 180°, penetrating deeper (8 Å) into a hydrophobic pocket. Disruption of the M794^7.32^-F780^6.57^-F776^6.53^ hydrophobic triad core and a conformational propagation path predominantly comprising TM6-7 residues are identified as key elements mediating force induced mGlu2 activation. Further *In vivo* rescue experiments support that mGlu2’s mechanosensitivity is dependent on FTM and is required for vestibular function. This work establishes a paradigm for class C GPCR mechanotransduction, revealing unprecedented structural mechanisms underlying force-induced Gq coupling and offering a chemical toolset to modulate mechanical signaling of GPCR.

## Introduction

The ability to perceive and respond to physical forces is a fundamental biological process happened across all domains of life^1–3^. Cells convert a wide range of mechanical stimuli, such as stretch, shear stress, and osmotic pressure, into intracellular electrical or chemical signals through a process known as mechanotransduction^4^. Among the known mechanosensitive proteins, ion channels exemplified by PIEZOs represent the most extensively characterized force sensors^5–7^. Their activation is primarily explained by two models: the “force-from-lipids” model, where changes in membrane tension and lipid packing gate the channel, and the “force-from-tether” model, where force is transmitted through physical linkages to the extracellular matrix or the cytoskeleton^7^. Beyond ion channels, emerging evidence indicates that various G protein-coupled receptors (GPCRs), the largest family of membrane receptors and premier drug targets, can be activated by forces and regulate critical processes such as cardiovascular regulation, balance perception and immunological functions ^8–10^. Intriguingly, force-induced activation of GPCRs, such as GPR68, LPHN2 and AT1R, often elicit downstream signaling pathways distinct from those activated by their canonical chemical ligands, suggesting divergent GPCR activation mechanisms between physical and chemical stimuli in selective physiological contexts ^8, 9, 11-14^. To date, the most clearly understood mechanism for GPCR mechanosensing is found in adhesion GPCRs (aGPCRs), which utilize a “*Stachel*” sequence that is exposed upon mechanical stimulation to activate the receptor^15–17^. Significant advances have also been made in identifying mechanosensitive receptors within non-aGPCRs, such as classes A and B members exemplified by S1PR1 and PTH1R; however, the molecular basis for their mechanosensitivity remains largely unresolved^18–21^. Notably, with different evolutionary route, it remains unclear whether the structurally distinct yet functionally important class C family of GPCRs also possesses intrinsic mechanosensitivity.

In our parallel work, we have identified a Class C GPCR, metabotropic glutamate receptor 2 (mGlu2), as a previously uncharacterized mechanosensor expressed in the kinocilium of vestibular hair cells, where it is essential for the perception of balance by activating the Gq-PLC-Ca^2+^ pathway. In the present study, leveraging a combination of structural, biochemical and cellular functional analyses, we revealed that the mechanosensitivity of mGlu2 is mediated by a unique tethered mechanism involving a N-terminal force transduction motif (FTM) that activates the receptor dimer in a *cis* manner. We then rationally developed an FTM-derived peptide that mimics force-induced mGlu2 activation. By resolving the Cryo-EM structures of mGlu2 at apo state, in complex with different peptides, as well as ternary complex of FTM bound mGlu2 and Gq trimer, we revealed an activation mechanism fundamentally different from the canonical activation induced by its endogenous agonist glutamate. The distinct structural mechanisms underlying force- or Glutamate-induced mGlu2 activation and downstream pathways were supported by our *in vivo* functional studies (Figure 1A). Our results provide the mechanistic framework for the mechanosensitivity of a Class C GPCR, special activation route and distinct G protein subtype coupling.

**Figure 1.**
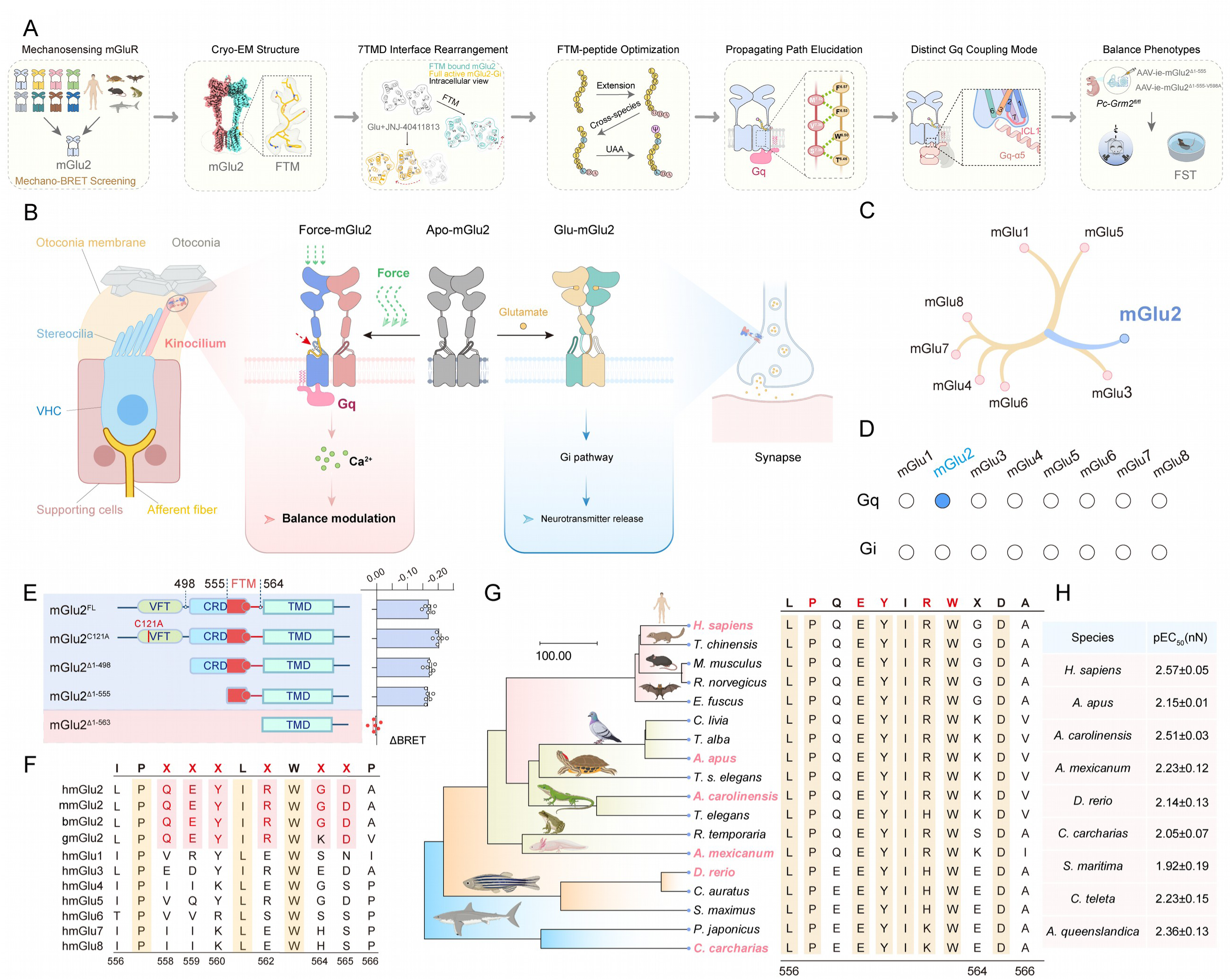
Mechanosensitivity of mGlu2 is regulated via a conserved N-terminal motif. **(A)** Schematic diagram illustrating the overall research flow for structural basis of asymmetric class C GPCR activation and distinct Gq coupling induced by force stimulation. **(B)** Schematic illustrating the mGlu2 activation in response to force or glutamate stimulation. mGlu2 expressed at the kinocilium of vestibular hair cell senses force and is essential for normal balance. Force stimulation activates the mGlu2-Gq pathway and regulates equilibrioception through Ca^2+^ signaling. Glutamate binds to synaptic mGlu2 and selectively activates Gi signaling to regulate neurotransmitter release. **(C)** Phylogenetic tree showing the evolutionary relationship of human mGlu receptors. **(D)** Summary of Gq and Gi signaling of human mGlu receptors in response to force stimulation. Note that only mGlu2 is sensitive to force stimulation, activating Gq signaling. See also Extended Data Figure 1 B-C. **(E)** Left panel: Schematic representation of different N-terminal truncated constructs of mGlu2, including full-length mGlu2 (referred to as mGlu2^FL^), mGlu2^Δ1-498^, mGlu2^Δ1-555^ and mGlu2^Δ1-563^. A C121A mutant was also generated based on mGlu2^FL^. Right panel: Force-induced Gq activation in HEK293 cells transfected with the different truncated constructs of mGlu2, as measured by G protein dissociation BRET assay. **(F)** Sequence alignment of the putative force transduction motif (FTM) of human mGlu2 with those from mGlu2 of other species or from other human mGluRs. Residues conserved in all the mGluRs (P557 and W563) are shown with a yellow background, and residues conserved only in mGlu2 from different species are colored red. **(G)** Sequence alignment of FTM in mGlu2 among different species of jawed vertebrates. Conserved residues are shown with a yellow background. The phylogenetic tree and the species divergence time are constructed using TimeTree (http://www.timetree.org/home). The animal images were generated using BioRender (https://biorender.com). **(H)** The presentation of pEC_50_ values for force-induced mGlu2 activation in various species. (E) Values are shown as the mean ± SEM of six independent experiments (n=6). \*\*\**P* < 0.001; ns, no significant difference, the mGlu2 mutation groups compared with the WT groups. The data were analyzed by one-way ANOVA with Dunnett’s multiple comparisons test.

## Results

### Force activates mGlu2 but not other 7 mGlu members through an N-terminal FTM motif

Our parallel study suggests that mGlu2 expressed in the kinocilium of vestibular hair cells is able to sense force, coupling to Gq and is essential for normal balance (Figure 1B). Several class A and class B GPCRs, such as GPR68 and PTH1R, are known to be activated by force^8, 16, 22, 23^. However, to our knowledge, no class C GPCR has been reported to be mechanosensitive. We therefore examined force-induced Gq activity of all 8 mammalian mGlu members using a magnetic beads assay^9^. Only mGlu2, but not the other mGlu members, is capable of initiating Gq signaling in response to mechanical stimulation (Figure 1C-D and Extended Data Figure 1B-C). This unique mechanotransduction property distinguishes mGlu2 from other mGlu family members.

Previous studies by our structural analysis and other chemical biology work have identified that the GPCR autoproteolysis-inducing (GAIN) domain of GPR133 and CD97 are primary force sensors for these two adhesion GPCRs^16, 24^. In detail, force engagement on the GAIN domain of these two receptors may cause dislodge of the *Stachel* sequences from the GAIN domain, following by the insertion of the *Stachel* sequences into the 7-helix transmembrane domain (7TMD) to induce receptor activation (Extended Data Figure 1A). Similar to adhesion GPCRs, the mGlu2 has very large N-terminal domain. We therefore hypothesized that certain fragment localized at the N-terminal domain of mGlu2, may serve as the hub of force sensor, which undergoes conformational change in response to force administration. We therefore constructed different truncations of N-terminal domain of mGlu2 according to its structural arrangement. The extracellular domain of mGlu2 is composed of a Venus flytrap (VFT) domain, which houses a positively-charged activation pocket, and a cysteine-rich domain (CRD). Initially, three truncations were designed, including the mGlu2^Δ1-498^ (lacking the VFT region), mGlu2^Δ1-555^ (lacking the VFT and most CRD region) and mGlu2^Δ1-563^ (lacking both VFT and CRD regions) (Figure 1E and Extended Data Figure 1D-E). Notably, removal of the first 555 residues (mGlu2^Δ1-498^ and mGlu2^Δ1-555^) did not significantly affect force-induced mGlu2 activation. In contrast, the deletion of residues 1-563 (mGlu2^Δ1-563^) completely eliminated the force-induced mGlu2 activation (Figure 1E). These findings indicate that the sequence spanning residues 556-563 is required for the mechanosensitivity of mGlu2. We therefore named this sequence the Force Transduction Motif (FTM) and designed the sequence spanning residues 556-566 as the Long Force Transduction Motif (LFTM).

We next conducted cross-species comparison of the putative LFTM sequences within the mGlu receptor family. Whereas all 11 residues of LFTM sequence of mGlu2 are highly conserved across mammalian species, other mGlu family members retain only 3 conserved amino acids (P^557^, I/L^561^ or W^563^), with the other residues showing significant sequence diversity (Figure 1F). This distinct evolutionary pattern may provide a molecular explanation for the exclusive mechanosensing capacity of mGlu2. Furthermore, phylogenetic analysis extends to non-mammalian vertebrates, including avian, amphibian, reptilian, and teleost species, and reveals remarkable conservation of mGlu2’s FTM sequence, showing that 8 out of 11 residues are mostly identical among different species. Importantly, all these mGlu2 orthologs from tested species are capable of sensing force to activate Gq signaling (Figure 1G-H and Extended Data Figure 1F-I). In addition, the residue at position 564 exhibits species-specific pattern, with glutamic acid (E) predominating in teleosts while lysine (K) in avian or amphibian orthologs (Figure 1G). This phylogenetic conservation of LFTM architecture suggests that mGlu2 may have maintained mechanosensitivity through evolutionary time (Figure 1G-H and Extended Data Figure 1F-I).

### FTM activated mGlu2 via a cis manner and development of mGlu2 peptide agonist

Because mGlu2 normally assumes dimer, the FTM may activate mGlu2 through two distinct modes, one interacting with the 7TM domain of the same protomer (Cis activation model), the other by interacting with the 7TM domain of the asymmetric subunit (Trans activation model) (Figure 2C). To investigate which interaction mode mediates force sensation by mGlu2, we constructed two mutant receptors: an FTM-Ala mutant (mGlu2^mFTM^) with mutations at the FTM and an ECL3-GSA mutant (mGlu2^mECL3^) with mutations in the 7TM domain (Figure 2A and Extended Data Figure 2A). The mGlu2^mFTM^ showed no Gq activity in response to force stimulation, whereas the mGlu2^mECL3^ displayed over 100-fold lower potency than WT mGlu2 (Figure 2B and Extended Data Figure 2B-C, 2E). We then co-expressed these two mutant receptors in the HEK293 cells and we reason that if mGlu2 could be activated by FTM via a trans mode, the heterodimer of mGlu2^mFTM^/mGlu2^mECL3^ should display enhanced mechanosensitity compared with either mutant (Extended Data Figure 2D). However, co-expression of mGlu2^mFTM^ with mGlu2^mECL3^ did not significantly increase the potency of force-stimulated Gq activation, which was comparable with that measure in HEK293 cells overexpressing mGlu2^mECL3^ alone (Figure 2B-C). Therefore, this observation supports a Cis activation mode, that is the two FTM sequences interact with their own 7TM domain to convey the force-induced mGlu2 activation.

**Figure 2.**
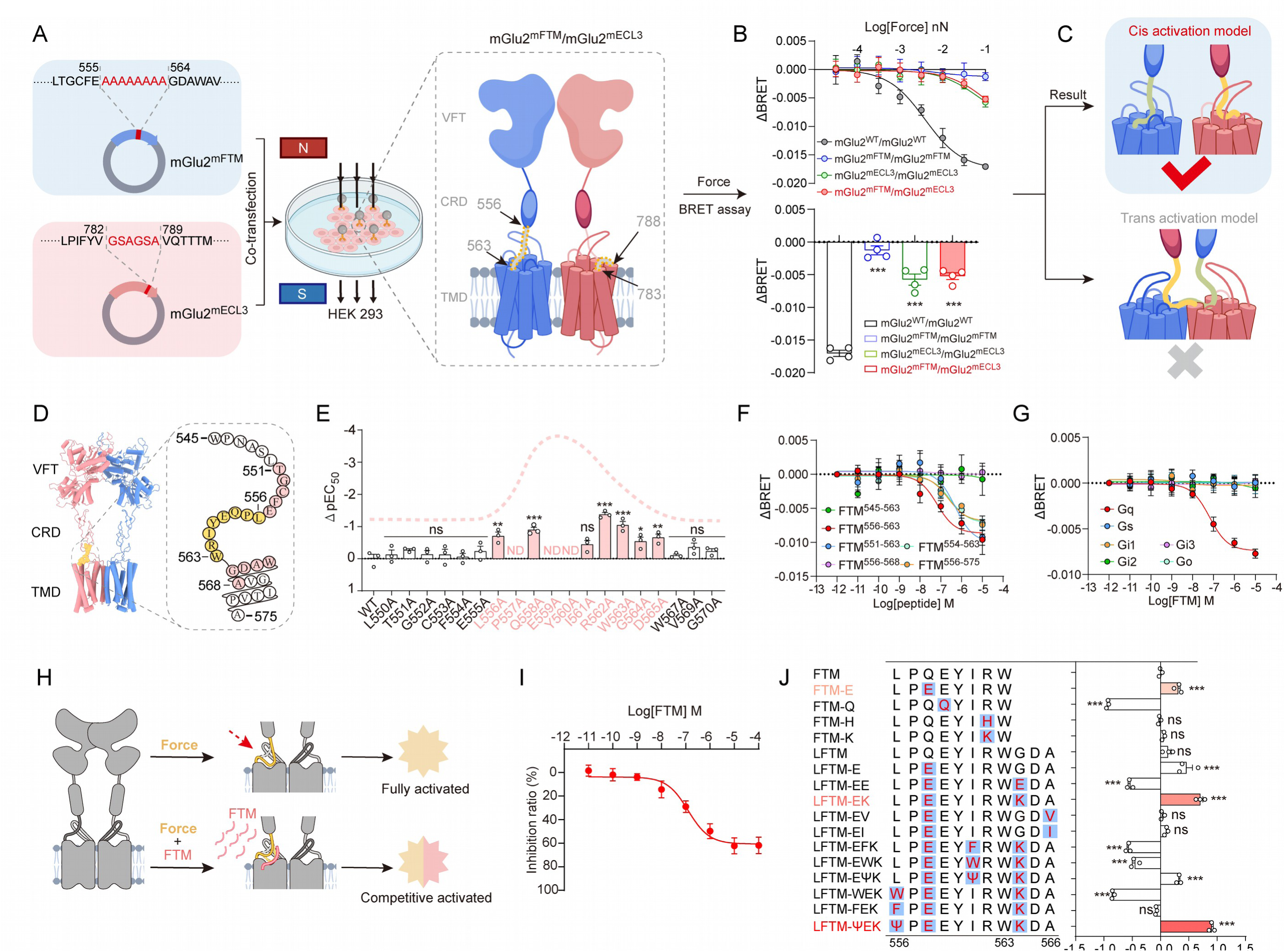
Force induces mGlu2 activation via a *cis* manner. **(A)** Schematic diagrams showing the strategy for investigating the activation mode of mGlu2 dimer. Two mGlu2 mutants, an FTM-deficient mGlu2^mFTM^ that was generated by replacing residues L556-W563 to Ala and an FTM-binding-deficient mGlu2^mECL3^ construct that was generated by replacing residues 783-788 with two GSA sequence, were employed to examine their effects on the mechanosensitivity of mGlu2 in HEK293 cells using BRET assay. The mGlu2^mFTM^ subunit was depicted in blue, and the mGlu2^mECL3^ subunit in red. Modified regions within both subunits were indicated by grey segments within yellow dashed boxes. **(B)** Dose-dependent Gq activation (upper panel) and the quantification of the maximal BRET value changes (lower panel) in mGlu2^WT^/mGlu2^mFTM^/mGlu2^mECL3^ hetero-/homodimers-overexpressed HEK293 cells in response to force applied via paramagnetic beads coated with anti-Flag M2 antibody. **(C)** Schematic of proposed cis or trans activation model of mGlu2 dimer. The cis activation model was highlighted with a blue background. Yellow and green segments represented the FTM in different subunits. **(D)** Structural schematic of the FTM region in mGlu2 (PDB ID: 7EPA). **(E)** Effects of mutations of residues within 550-570 on the force-induced Gq activation in mGlu2-overexpressed HEK293 cells. **(F)** Dose-dependent Gq activation in mGlu2-overexpressed HEK293 cells in response to stimulation with different synthetic peptides derived from FTM sequence. Representative curve data are shown as mean ± SEM from three independent experiments (n=3). **(G)** Dose-dependent Gq/Gs/Gi/Go activation in mGlu2-overexpressed HEK293 cells in response to FTM peptide stimulation. Representative curve data are shown as mean ± SEM from three independent experiments (n=3). **(H)** Schematic of the competition assay between FTM peptide and mechanical force for mGlu2 activation. mGlu2 can be fully activated by force, involving conformational changes in the segment L556–W563. Stimulation with FTM peptide competes with force-mediated activation, leading to reduced force-induced activation with increasing FTM concentration. Yellow stars indicate the level of force-induced activation; pink stars represent FTM peptide-induced activation. The yellow curve shows the position of the FTM motif in mGlu2 under force stimulation, with or without FTM peptide. The grey dashed curve indicates the position of the FTM motif in the apo state. The pink curve represents the FTM peptide. **(I)** Representative competitive binding curves of FTM peptide stimulation versus force stimulation on mGlu2 activation. Representative curve data are shown as mean ± SEM from three independent experiments (n=3). **(J)** Summary of the potencies of different FTM-derived peptides in activating the Gq signaling in mGlu2-overexpressed HEK293 cells. The sequence 556-566 in mGlu2 is designated as LFTM, with the symbol Ψ representing trimethylphenylalanine. Bar graph showed the difference in potency (ΔpEC_50_) between FTM and FTM peptide variants, generated from data shown in Extended Data Figure 2K-2N. Mutated residues are highlighted in red on a blue background. Peptides and corresponding bars showing significantly improved activation potency are colored light pink to red. (B, E, J) Values are shown as the mean ± SEM of three independent experiments (n=3). (B) \*\*\**P* < 0.001, the mGlu2^mutation^/mGlu2^mutation^ groups compared with the mGlu2^WT^/mGlu2^WT^ groups; (E)\**P*<0.05; \*\**P*<0.01; \*\*\**P* < 0.001; ns, no significant difference; ND, not detected, the mGlu2 mutation groups compared with the WT groups; (J) \**P*<0.05; \*\**P*<0.01; \*\*\**P* < 0.001; ns, no significant difference; ND, not detected, the FTM mutation groups compared with the FTM groups. All the data were analyzed by one-way ANOVA with Dunnett’s multiple comparisons test.

To further explore the mechanism underlying the FTM-mediated mGlu2 activation, we performed mutational analysis of FTM sequence (Figure 2D). We revealed that mutations of key hydrophobic or charge residues in FTM motif, such as P557, E559, Y560, R562 and W563, to alanine significantly impaired the force-induced mGlu2 activation. Importantly, the P^557^XE^559^YXRW^563^ motif is highly conserved across different species, supporting an important role of the FTM sequence in mGlu2’s response to mechanical stimuli (Figure 2E and Extended Data Figure 2F-J). We hypothesized that a synthetic peptide corresponding to FTM sequence may directly activate mGlu2 by mimicking the force administration. Consistently, synthetic FTM peptides, such as FTM^556–563^ and FTM^556–575^, only induced Gq signaling, but not other G protein pathways (Figure 2F-G). Intriguingly, the FTM^556–563^ peptide dose-dependently compete with force-induced mGlu2 activation (Figure 2H-I), further suggesting that the FTM is the hub in mediating force-induced mGlu2 activation. Although most of the FTM sequence is highly conserved among different species, variations were observed at positions 558 and 562, primarily in fish species (Figure 1G). By testing different sequence mutations, we found that Q558E-G564K double mutant of LFTM exhibited an enhanced potency to activate Gq, with an EC_50_ of 10.9±1.4 nM (Figure 2J and Extended Data Figure 2K-L). Moreover, by introducing the non-natural amino acid trimethylphenylalanine at position 556 (L^556^TMF), which is subsequently denoted by the symbol Ψ, the potency of mGlu2-Gq activation was further increased by approximately 1.3-fold, with an EC_50_ value of 8.6±1.6 nM (Figure 2J and Extended Data Figure 2M-N).

### Overall structure of mGlu2 in complex with FTM peptide and the Gq trimer

To understand the structural basis of mGlu2 activation by FTM, which may mimic the force-induced mGlu2 activation, we reconstituted the FTM-mGlu2 complexes *in vitro* and performed structural analysis via Cryo-EM. Two different mGlu2 dimer conformations were identified during the 3D classification, which resulted in the structures of the apo-mGlu2 dimer (3.4 Å) and the FTM-mGlu2 dimer (3.4 Å) (Figure 3A and Extended Data Figure 3). To further stabilize the mGlu2-Gq complex, we employed the more potent LFTM-ΨEK peptide agonist. Classification and structural calculations resulted in structures of the LFTM-ΨEK-mGlu2 dimer (2.9 Å) and the LFTM-ΨEK-mGlu2-Gsq complex (6.20 Å) (Figure 3A and Extended Data Figure 4-5). The structure of the LFTM-ΨEK-mGlu2 (19-857) dimer closely resembles that of the FTM-mGlu2 (19-872) dimer, with an overall root mean square difference (RMSD) of 0.37 Å (Figure 3A and Extended Data Figure 6A, 9M-N).

**Figure 3.**
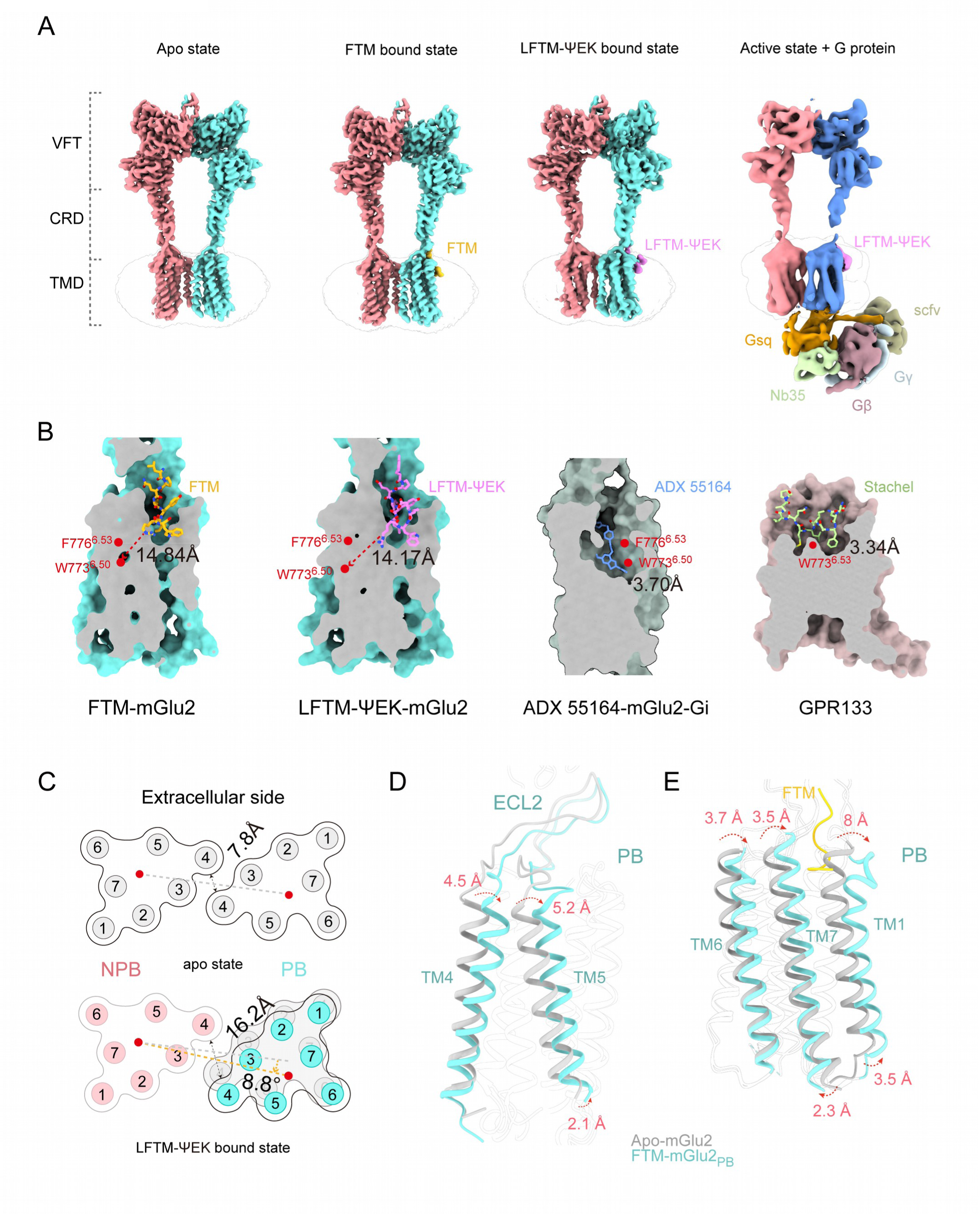
Overall structures of mGlu2 in different states. **(A)** Orthogonal views of the density map for the apo-mGlu2, FTM-mGlu2 dimer, LFTM-ΨEK-mGlu2 dimer, and the LFTM-ΨEK-mGlu2-Gsq complexes. The NPB subunit is shown in red, the PB subunit in teal or blue. FTM is colored gold; LFTM-ΨEK is shown in purple; Gsα, Gβ, Gγ, scFv16, and Nb35 are depicted in orange, reddish brown, grayish blue, grayish green, and grass green, respectively. **(B)** Cut-away view of the ligand-binding pocket in the FTM-mGlu2 dimer complex, LFTM-ΨEK-mGlu2 dimer complex, ADX 55164-mGlu2-Gi complex (PDB ID: 7MTR) and GPR133(PDB ID: 7EPT). FTM, LFTM-ΨEK, ADX 55164, and the GPR133 *Stachel* motif are shown as yellow, purple, blue, and green sticks, respectively. The distance between the ligand and the toggle switch position are shown as dashed lines. **(C)** Extracellular view of the dimerization mode of the TMD during mGlu2 receptor activation. apo-mGlu2 is shown in grey; LFTM-ΨEK-mGlu2 dimer is shown in pink (NPB subunit) and teal (PB subunit). Subunit centers are marked by red dots connected by a dashed line. Distance between TM4 helices is indicated by a black double-headed arrow. The yellow dashed arrow indicates the global rotation angle of the TMD from the apo-mGlu2 conformation to the LFTM-ΨEK-mGlu2 dimer conformation. **(D-E)** Comparison between apo-mGlu2 and PB protomer of FTM-mGlu2 dimer structures focusing on TM4-TM5 (D) and TM1-TM6-TM7 (E). mGlu2 in FTM-mGlu2 dimer and apo-mGlu2 structures is shown in teal and grey, respectively. Red dashed arrows indicate the direction of movement.

One intriguing discovery is that only one FTM or LFTM-ΨEK peptide could be assigned in one protomer in the dimer of FTM-mGlu2, LFTM-ΨEK-mGlu2 or LFTM-ΨEK-mGlu2-Gsq complex structures (Figure 3A). Notably, the Gsq only binds to peptide-bound (PB) protomer of mGlu2, but not the non-peptide-bound (NPB) protomer (Figure 3A). The LFTM-ΨEK bound position and the peptide-induced Gsq coupling to the same mGlu2 protomer are in consistent with the cis activation model of mGlu2 in response to force stimulation (Figure 2C, 3A). Despite the selective binding of the peptide to only one protomer, the two protomers of mGlu2 show very similar structures, as indicated by a TMD RMSD value of only 1.8 Å (Extended Data Figure 6D). In contrast, the two protomers of FTM (mGlu2^FTM^) or LFTM-ΨEK bound mGlu2 (mGlu2^LFTM-ΨEK^) showed larger structural differences compared with the mGlu2^apo^ (Extended Data Figure 6B -C). The RMSD between the PB protomers or NPB protomers of the mGlu2^FTM^ and mGlu2^apo^ is 2.3 Å and 2.0 Å, respectively; with the largest difference identified at the extracellular ends of TM1, ICL1 and ICL2 for PB protomer and ICL2 for NPB protomer (Extended Data Figure 6B).

### An atypical binding position of the FTM peptide

The FTM peptide binds within the 7TM domain, but not the orthosteric site located in the VFT domain encompassing the endogenous agonist glutamate^25^ (Figure 3A). Contrary to the deep insertions of the positive allosteric modulator (PAM) ADX 55164 into TM5 and TM6 in the mGlu2-Gi complex, FTM and LFTM-ΨEK bind to mGlu2 at a shallower location within a pocket formed between TM1, TM7 and ECL2, as indicated by long distance between the FTM binding pocket and the W773^6.50^XXF^6.53^ motif (Figure 3B). It’s worth noting that the direct contact between the *Stachel* sequence of aGPCRs and the “toggle switch” W^6.53^ serves as the key mechanism in force-induced activation of the aGPCRs, such as GPR133^16^ (Figure 3B). Therefore, the peptide binding mode and force sensation mechanism of mGlu2 may be distinct from those of mechanosensitive aGPCRs.

### The 7TM dimer interface

In the mGlu2^apo^ structure, the TM4 of the two protomers are in close proximity, as previously reported ^26^. However, the FTM-mGlu2 and LFTM-ΨEK-mGlu2 dimers do not exhibit the torsion of 7TMD observed in the previously reported intermediate state of mGlu2 in complex with its endogenous agonist glutamate. The FTM-mGlu2 and LFTM-ΨEK-mGlu2 dimers exhibit striking similarities in the overall packing arrangements of their transmembrane helices. Specifically, both dimers adopt an asymmetric 7TM dimerization interface mediated by interactions between TM4 of the NPB protomer and TM3 of the PB protomer (Figure 3C and Extended Data Figure 6A). Notably, this structural organization is significantly different from the TM4-centered symmetric 7TM dimer interface observed in the mGlu2^apo^, or the TM6-driven asymmetric interface configuration identified in the glutamate-bound mGlu2-Gi complex (Figure 3C, 4A and Extended Data Figure 6A).

**Figure 4.**
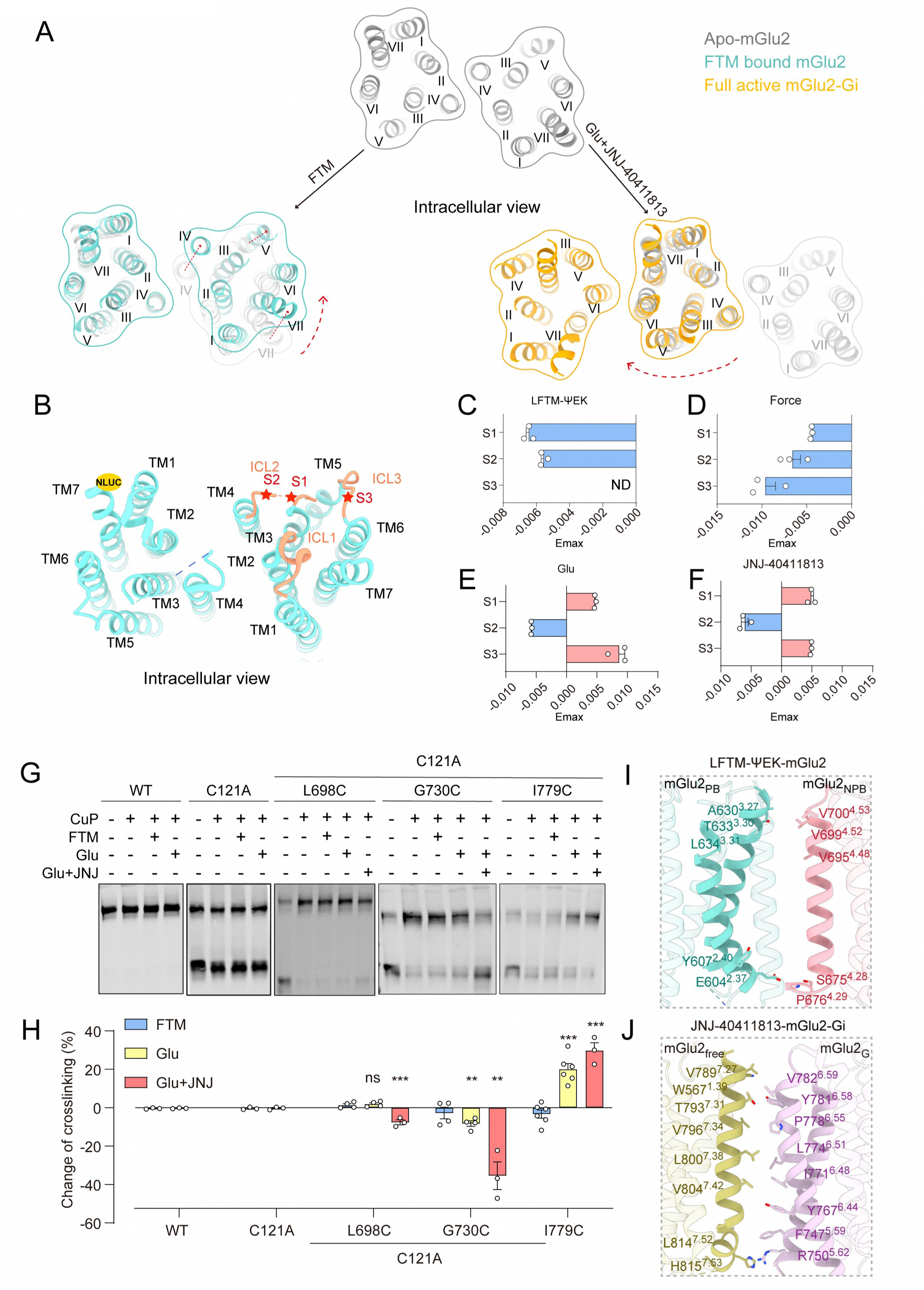
Dimerization interface of force-induced mGlu2 activation. **(A)** Multiple dimerization conformations of the TMD observed from the intracellular side during activation of the mGlu2 receptor bound to different agonists. Red dashed arrows indicate conformational changes in the receptor TMD from the apo state to the activated state upon agonist binding. apo-mGlu2, FTM-mGlu2 dimer, and Glu-JNJ40411813-mGlu2-Gi (PDB ID: 7E9G) complex are depicted in grey, cyan, and yellow, respectively. **(B)** Schematic representation of the FlAsH-BRET assay design. The NLUC was inserted at the C-terminal of mGlu2, and the FlAsH motif were inserted in designated positions in the other protomer (as shown in the figure). In response to binding of agonists (LFTM-ΨEK, Glu and JNJ-40411813) or antagonists (LY341495), the ICL2-S1/2 and the ICL3-S3 moved close to or away from the C-terminus of mGlu2, respectively. **(C-F)** The maximal response of four FlAsH–bioluminescence resonance energy transfer biosensors of mGlu2 in response to LFTM-ΨEK(C), force(D), Glu(E) and JNJ-40411813(F) stimulation. The values are the mean ± SEM of three independent experiments (n = 3). **(G)** Cross-linking analysis of cell surface Snap-tagged mGlu2 WT or mutants containing cysteine substitutions in TM4, TM5 or TM6. Cells were treated with (+) or without (−) with FTM, Glu, Glu+JNJ-40411813 and CuP. Snap-mGlu2 monomers and dimers were separated in SDS-PAGE and detected via the fluorophore covalently attached to the receptors. **(H)** The percentage of dimers relative to the total amount of mGlu2 was quantified by imaging the fluorescent blots in (G). **(I-J)** Interactions between the mGlu2_PB_ and mGlu2_NPB_ protomers in the LFTM-ΨEK-mGlu2 dimer (I) and JNJ-40411813-mGlu2–Gi structures (J). The residues that are involved in dimer interactions or in close proximity are shown as sticks. In the LFTM-ΨEK-mGlu2 dimer complex, the mGlu2_PB_ and mGlu2_NPB_ protomers are colored teal and red, respectively. In the JNJ-40411813–mGlu2–Gi complex, the G protein-bound subunit (mGlu2_G_) and the G protein-free subunit (mGlu2_free_) of mGlu2 are shown in purple and brown, respectively. **(H)** Values are shown as the mean ± SEM of at least three independent experiments (n=3). \*\**P*<0.01; \*\*\**P* < 0.001; ns, no significant difference, all treatment groups compared with FTM treatment group. The data were analyzed using Ordinary one-way ANOVA followed by Tukey’s multiple comparisons test.

From the extracellular side view, the binding of FTM or LFTM-ΨEK peptide caused anti-clockwise rotation of 7TM bundles of NPB protomer by 8.8°, using NPB protomer as a reference. These rotations moved the TM4 away from two protomers by approximately 8.4 Å, thus disrupting previous interface (Figure 3C). The binding of FTM peptide and its unique rotational mechanism induced significant helical displacements in the TMD regions of both PB and NPB protomers of mGlu2^FTM^. In the PB protomer, peptide binding primarily drove outward counterclockwise rotation at the extracellular termini of TM1 and TM7, with displacements of 8.0 Å and 3.5 Å compared with mGlu2^apo^, respectively (Figure 3E). The extracellular end of TM6 coordinated a 3.7 Å displacement following TM7’s movement (Figure 3E). Importantly, these extracellular rearrangements were accompanied by a 2.3 Å and 3.5 Å outward displacement of the intracellular termini of TM7 and TM1, respectively (Figure 3E and Extended Data Figure 6E). These movements potentially expose structural motifs critical for forming the shallow groove required for Gq protein coupling.

### A non-classical 7TM rearrangement of mGlu2

Previous studies have suggested that the relative movement of the two protomers within the class C GPCR dimer is the key event in the transition from inactive to active state, as exampled by the comparison between the structures of mGlu2^apo^ and the glutamate-JNJ-40411813-mGlu2-Gi complex^25, 27–29^. Specifically, in the apo state, the two protomers of the mGlu2 are mainly stabilized by the TM3-TM4 interface. After binding to glutamate and JNJ-40411813, the 7TM rearranges to form a stable structure mainly maintained by the TM6-TM6 interaction (Figure 4A). In contrast, mGlu2^FTM^ exhibits a distinct conformational landscape, characterized by an 8.8° rotation of the PB protomer (Figure 3C).

To probe LFTM-ΨEK-induced conformational changes in mGlu2, we employed an intramolecular FlAsH-Bioluminescence Resonance Energy Transfer (FlAsH-BRET) assay ^30–34^. We attached the Nluc at the C-terminus of one mGlu2 protomer and inserted the FlAsH probes at the intracellular loops of the other protomer (Figure 4B and Extended Data Figure 7A). In HEK293 cells overexpressing these FlAsH-BRET sensors, quantitative analysis revealed that glutamate, PAM (JNJ-40411813), antagonist (LY-341495), LFTM-ΨEK and force all induced a decrease of BRET signal at the S2 position (TM4-terminus of ICL2), indicating a dissociation between this site and the C-terminus of the other protomer (Figure 4C-F and Extended Data Figure 7B-E). The FlAsH probes at the other sites revealed divergent movements of the intracellular loops of mGlu2 dimer in response to different ligands stimulation. For example, whereas the LFTM-ΨEK stimulation decreases the BRET signal at the S1 position (TM3-terminus of ICL2), the other ligands exerted an opposite effect (Figure 4C-F and Extended Data Figure 7B-E). Moreover, while both glutamate and JNJ-40411813 induced an assembly between the S3 site at ICL3 (between TM5 and TM6) and the Nluc-tagged C-terminus of the other protomer, LFTM-ΨEK application did not show detectable effects (Figure 4C-F and Extended Data Figure 7B-E). Remarkably, mechanical force stimulation replicated LFTM-ΨEK-induced BRET change patterns, suggesting that the FTM-derived peptide recapitulates force-induced activation (Figure 4C-D and Extended Data Figure 7C-D). Collectively, these data suggest divergent activation mechanisms of mGlu2 in response to physical and chemical stimuli.

We next performed the classical disulfide crosslinking experiments to examine mGlu2 dimer interfaces in response to LFTM-ΨEK or glutamate stimulation, which have been extensively used for the characterization of class C GPCR activation ^35, 36^. Using a cysteine-depleted mutant (mGlu2-C121A) with an N-terminal SNAP tag for selective labeling, we observed that WT mGlu2 predominantly formed dimers under non-reducing conditions, whereas mGlu2-C121A presented as monomers (Figure 4G). Glutamate stimulation reduced TM4-TM4 (L698^4.51^) and TM5-TM5 (G730^5.42^) crosslinking while enhancing TM6-TM6 (I779^6.56^) interactions, consistent with the observed TM6-TM6 interaction in the fully activated mGlu2 bound with glutamate and JNJ-40411813 (Figure 4G-H). In contrast, FTM peptide stimulation did not induce significant crosslinking at TM3-TM3 (T633^3.30^, L634^3.31^), TM4-TM4 (I674^4.27^, S675^4.28^, P676^4.29^, S678^4.31^, L698^4.51^, E701^4.54^), TM5-TM5 (G730^5.42^, Y734^5.46^), or TM6-TM6 (I779^6.56^) (Figure 4G-H and Extended Data Figure 7F-G). These specific changes detected by disulfide cross-linking align well with the structural observation and the result from FlAsH-BRET assay, indicating a novel mechanism of mGlu2 activation induced by FTM, which recapitulates force application. Supporting this hypothesis, mutations of the residues at the mGlu2 dimer interface that are essential for glutamate-induced mGlu2-Gi signaling showed minimal effects on LFTM-ΨEK-induced mGlu2-Gq activation (Figure 4I-J and Extended Data Figure 7H-K).

### Detailed interactions between FTM and mGlu2

Both FTM and LFTM-ΨEK exhibited extended β-sheet structures, binding between the CRD, TM1, TM7 and ECL2-3 regions of mGlu2 (Figure 5A-B, 5D and 6A-B). In FTM-mGlu2 complex, the Q558^FTM^ engaged in hydrogen bonds with E708^ECL2^ and Y787^ECL3^, whereas the E559 ^FTM^ formed charge-charge interaction with R720^ECL2^ and hydrogen bond with T718^ECL2^ (Figure 5C, 5F-G and Extended Data Figure 8A-B). The N-terminal residues L556^FTM^ formed hydrophobic or van der wal interactions with R714^ECL2^, E715^ECL2^, V717^ECL2^ of ECL2 (Figure 5E and Extended Data Figure 8A-B). At the C-terminus of the peptide, the I561^FTM^ formed hydrophobic or van der Waals interactions with I624^2.57^, C795^7.33^ and V796^7.34^ (Figure 5H and Extended Data Figure 8A-B). Consistent with these observations, mutations of L719^ECL2^, V717^ECL2^ and R714^ECL2^ to alanine significantly diminished the peptide-induced mGlu2-Gq activation (Figure 5I and Extended Data Figure 8D, 8F). Notably, the mutational effects on these pocket residues were similar between force-induced and FTM peptide-induced mGlu2-Gq activation, further supporting the FTM-regulated mechansensitive mechanism of mGlu2 (Figure 5I-J and Extended Data Figure 8D-F).

**Figure 5.**
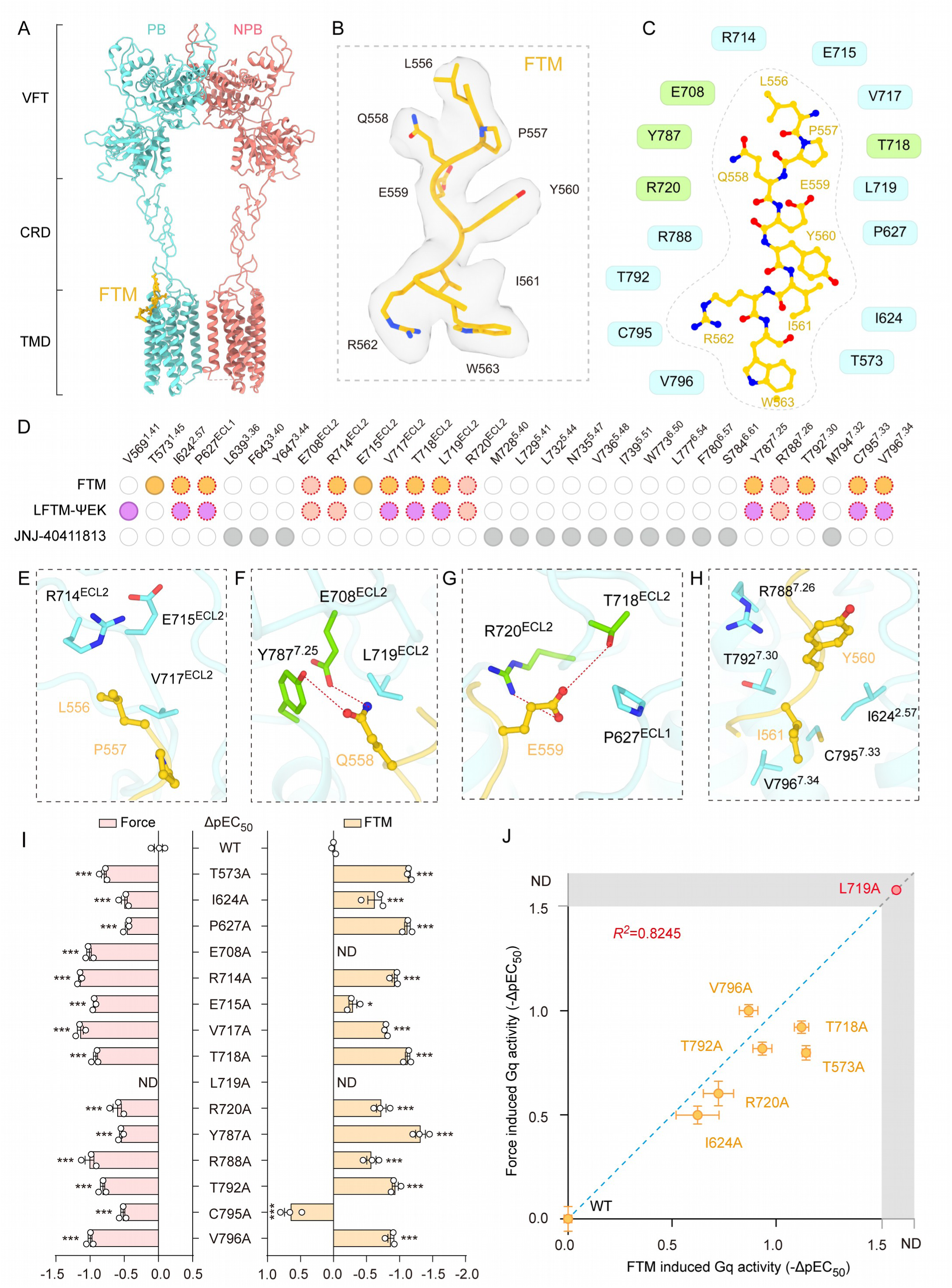
FTM binding mode and detailed interaction. **(A)** Cryo-EM structure of the FTM-mGlu2 dimer complex. The PB and NPB protomers of mGlu2 are coloured Turquoise and red, respectively. The FTM peptide is colored in yellow. **(B)** Cryo-EM density of FTM peptide. FTM is colored in yellow, and the EM density is shown in grey. **(C)** Schematic of interactions between mGlu2 residues and ago-PAM FTM peptide. Lawn Green, hydrogen bond or charge interaction; Turquoise, hydrophobic interaction. **(D)** Comparison of the FTM, LFTM-ΨEK or JNJ-40411813 binding pocket in mGlu2. The residues that engage with FTM, LFTM-ΨEK and JNJ-40411813 are shown in Amber Glow, purple and gray, respectively. The residues forming hydrogen bonds or electrostatic interactions with FTM or LFTM-ΨEK are depicted in shell pink, specially. The residues, which interacts with both FTM and LFTM-ΨEK, are shown in red dashed borders. **(E-H)** Detailed representation of the binding pockets involving residues L556-P557 (E), Q558 (F), E559 (G), and Y560-I561 (H) on the FTM peptide. Lawn Green, hydrogen bond or charge interaction; Turquoise, hydrophobic interaction. **(I)** Effects of alanine mutations within the FTM binding pockets of mGlu2 on FTM- and Force-induced Gαq-Gγ dissociation in mGlu2-overexpressing cells. **(J)** Summary of the effects of mutations of FTM-interacting pocket residues on the FTM- and force-induced Gq activity measured by BRET assay. The X axis represents -ΔpEC_50_ values between FTM-induced Gq activity of mGlu2 mutants and mGlu2-WT; positive values indicate reduced potency of FTM. The Y-axis represents -ΔpEC_50_ values between force-induced Gq activity of mGlu2 mutants and mGlu2-WT; positive values indicate reduced potency of force. Mutants in the first quadrant displayed both impaired FTM response and loss of force sensitivity, mimicking the coordinated changes in FTM- and force-induced mGlu2 Gq activity modulation (n=3). The blue dashed line represents the y=x reference, while the R² value reflects the correlation between mutant-induced differences in FTM- and force-stimulated Gq activity changes. (I) Values are shown as the mean ± SEM of three independent experiments (n=3). \**P*<0.05; \*\*\**P*<0,001; ND, not detected, the different mutations of mGlu2 compared with the WT group. The data were analyzed by one-way ANOVA with Dunnett’s multiple comparisons test.

**Figure 6.**
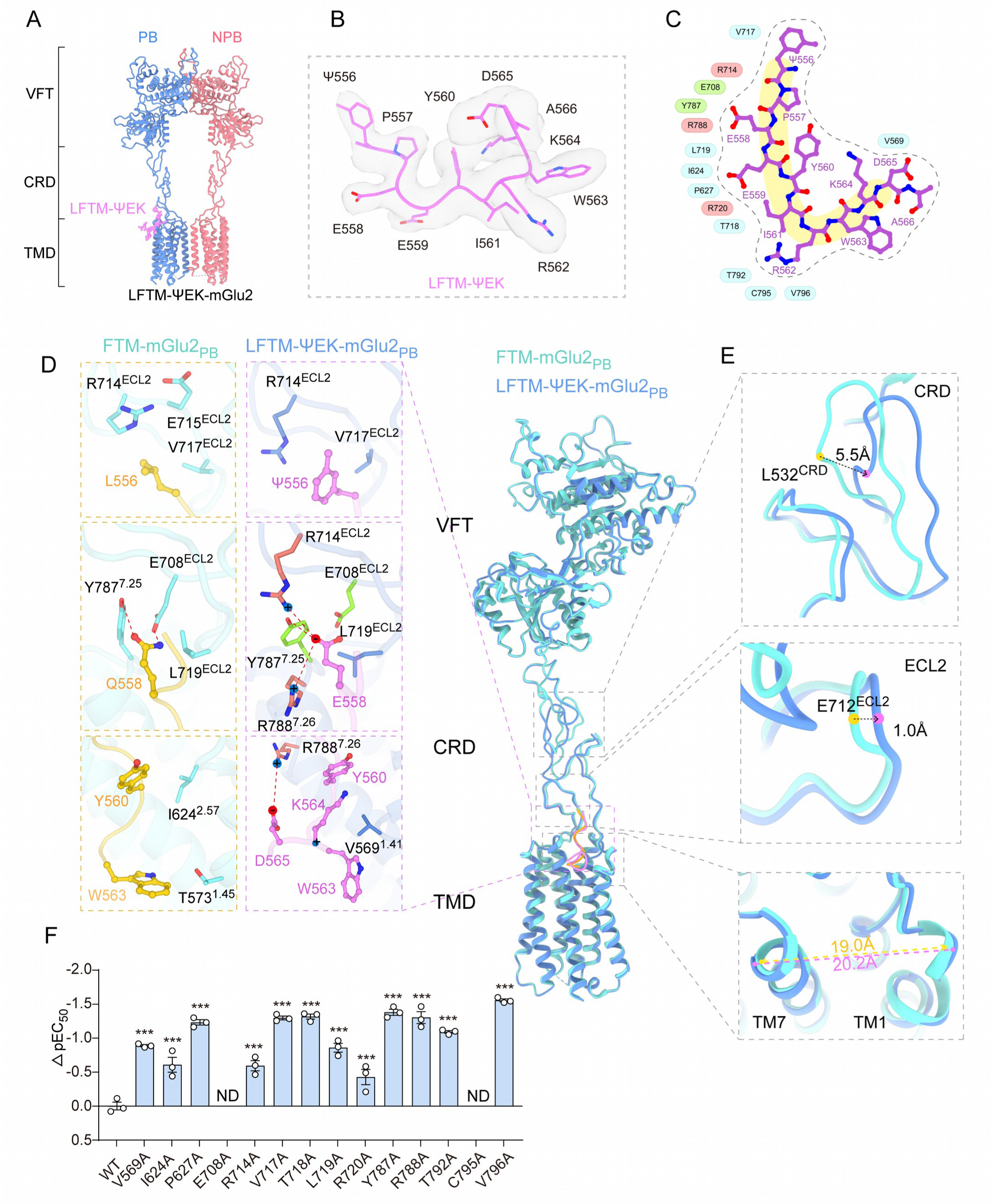
LFTM-ΨEK interaction with mGlu2. **(A)** Cryo-EM structure of the LFTM-ΨEK -mGlu2 complex. The PB and NPB protomers of mGlu2 are coloured blue and red, respectively. The LFTM-ΨEK peptide is colored in purple. **(B)** Cryo-EM density of LFTM-ΨEK peptide. LFTM-ΨEK is colored in purple, and the EM density is shown in grey. **(C)** Schematic of interactions between mGlu2 residues and ago-PAM LFTM-ΨEK peptide. Lawn Green, hydrogen bond; red, charge interaction; Turquoise, hydrophobic interaction. **(D-E)** The PB protomers of the LFTM-ΨEK–mGlu2 dimer and FTM–mGlu2 dimer complexes were aligned along TM2 and TM3. (LFTM-ΨEK-mGlu2_PB_, blue; FTM -mGlu2_PB_, Turquoise; FTM peptide, Amber Glow; LFTM-ΨEK peptide, purple). (D)Detailed comparative representation of the binding pockets of key residues at identical positions on FTM and LFTM-ΨEK. Relative to FTM, key residues mediating newly formed hydrogen-bonding and charge-charge interactions in the LFTM-ΨEK binding pocket are colored Lawn Green and red, respectively. (E) Detailed representation of discordant structural features between LFTM-ΨEK and FTM subunits. Residue positions in FTM -mGlu2_PB_ and LFTM-ΨEK-mGlu2_PB_ are indicated by yellow and purple dots, respectively; their distance is represented by a black dashed arrow. The distances between TM1 and TM7 in the two structures are shown with yellow and purple dashed lines, respectively. **(F)** Effects of different mutations within the binding pockets of mGlu2 on LFTM-ΨEK-induced Gαq-Gγ dissociation in mGlu2 overexpressing cells. (F) Values are shown as the mean ± SEM of three independent experiments (n=3). \*\*\**P* < 0.001; ND, not detected, the mGlu2 mutation groups compared with the WT groups. All the data were analyzed by one-way ANOVA with Dunnett’s multiple comparisons test.

### Comparison between LFTM-ΨEK- and FTM-induced mGlu2 activation

Compared with the FTM peptide, the modified LFTM-ΨEK peptide shares the similar pocket location but assumes an overall “fishhook” shape and steps closer to the toggle switch W773^6.50^ (Figure 3B, 6A-C and Extended Data Figure 9E). Since the side chains of R720^ECL2^, Y787^7.25^ and R788^7.26^ along the pocket of mGlu2 structure couldn’t be assigned into EM density, we performed molecular dynamics simulation to position their configurations (Extended Data Figure 9H-9J).

Specifically, the substitution of L^FTM^ to Ψ^LFTM-ΨEK^ at position 556 at the N-terminus of the peptide improved its rigidity. The side chain of Ψ556^LFTM-ΨEK^ rotated by approximately 90 degree compared with the L^FTM^, and formed potential cation: π interaction with R714^ECL2^ (Figure 6D and Extended Data Figure 9E). The rotation of Ψ556^LFTM-ΨEK^ also enabled a closer contact between the P557^LFTM-ΨEK^ and Y560^LFTM-ΨEK^ inside the peptide (Extended Data Figure 9E). The replacement of Q^FTM^ with E^LFTM-ΨEK^ at position 558 adds new charge interactions with R714^ECL2^ and R788^7.26^, allowing the peptide to penetrate deeper into the ligand pocket and to form a potential hydrogen bond with Y787^7.25^ (Figure 6D and Extended Data Figure 9A-D). At the C-terminus, the K564^LFTM-ΨEK^ engaged a cation-π interactions with Y560^LFTM-ΨEK^ within LFTM-ΨEK, which enabled an overall “fishhook” configuration of the peptide. The D565^LFTM-ΨEK^ form charge-charge interactions with R788^7.26^ (Figure 6D and Extended Data Figure 9E). All the above interactions between LFTM-ΨEK and mGlu2 are supported by mutational analysis, and these interactions may contribute to the higher affinity of LFTM-ΨEK over FTM in binding to mGlu2 (Figure 6F and Extended Data Figure 9E-G, 9K). Compared to the FTM-mGlu2 complex, binding of LFTM-ΨEK induce deflections of 5.5 Å at CRD, 1.0 Å at ECL2, and 1.2 Å between TM1 and TM7 in the PB protomer (Figure 6E and Extended Data Figure 9L-N). These observations suggest the structural plasticity of the FTM binding pocket, which may facilitate allosteric agonist design targeting mGlu2.

### Differential Coupling Mechanisms of mGlu2 with Gi and Gq

The LFTM-ΨEK-mGlu2-Gq complex structure has a relatively low resolution, with that at the interface between 7TMDs of mGlu2 and the Helix 5 of Gαq reaching 5.7 Å (Extended Data Figure 5). However, continuous densities for these regions could be traced, which enable easily main-chain tracing of the helices. Specifically, the side-chain densities of several key residues—W567^1.39^, R591^1.63^, L610^2.43^, L650^3.47^, I682^4.35^, L684^4.37^, L686^4.39^, F747^5.59^, F761^6.38^, F764^6.41^and F776^6.53^ in the TMD of PB protomer, W567^1.39^, F584^1.56^, L744^5.56^, F747^5.59^, R750^5.62^, K760^6.37^, W767^6.44^ and F810^7.48^ in the TMD of NPB protomer, as well as N392^G.H5.24^, L393 ^G.H5.25^ and V394 ^G.H5.26^ within the GαH5 domain—were clearly resolved, which helped determine the orientation of the helical main chains and assignment of residue numbers (Extended Data Figure 10A). Furthermore, computational simulations supported our structural model of the interaction mode between mGlu2 and Gq (Extended Data Figure 10B). Distinct from the canonical mGlu2-Gi complex, the LFTM-ΨEK-mGlu2-Gq complex preserves the TM3/4-mediated dimerization interface observed in LFTM-ΨEK-mGlu2 complex, eschewing the TM6-driven asymmetric dimer configuration (Figure 7A-B). Compared with the Gi in the glutamate-bound mGlu2-Gi complex, which selectively engages ICL2/ICL3-TM3 domains with its α5 helix projecting toward the TM4-TM2 interhelical cleft, the Gq that couples with LFTM-ΨEK-mGlu2 occupies a distinct pocket formed by ICL1-TM3/6/7, its α5 helix oriented orthogonally toward TM6 (Figure 7A, 7F and Extended Data Figure 10C). These distinct binding modes result in a 115° spatial angle between the α5 helices of Gi and Gq, projecting near-anti-parallel orientations in the membrane plane (Figure 7A and Extended Data Figure 10D). Compared to the Gi-bound state, when viewed from the intracellular perspective, TM4-TM6 of the Gq-bound mGlu2 exhibit clockwise displacements at their intracellular ends (Extended Data Figure 10E). TM4 displays a 5.2 Å outward displacement relative to the membrane plane, accompanied by 3.0 Å and 4.0 Å shifts in TM5 and TM6, respectively (Figure 7C and Extended Data Figure 10E). In contrast, the displacement of the extracellular ends of these TMs are relatively smaller (TM4: 3.7 Å, TM5: 1.7 Å, TM6: 0 Å) (Figure 7C). These helical movements likely expand the intracellular cavity of the 7TM bundle in the LFTM-ΨEK-mGlu2-Gsq complex, enabling the penetration of α5 helix of Gq approximately 8.3 Å deeper than that of Gi (Figure 7A).

**Figure 7.**
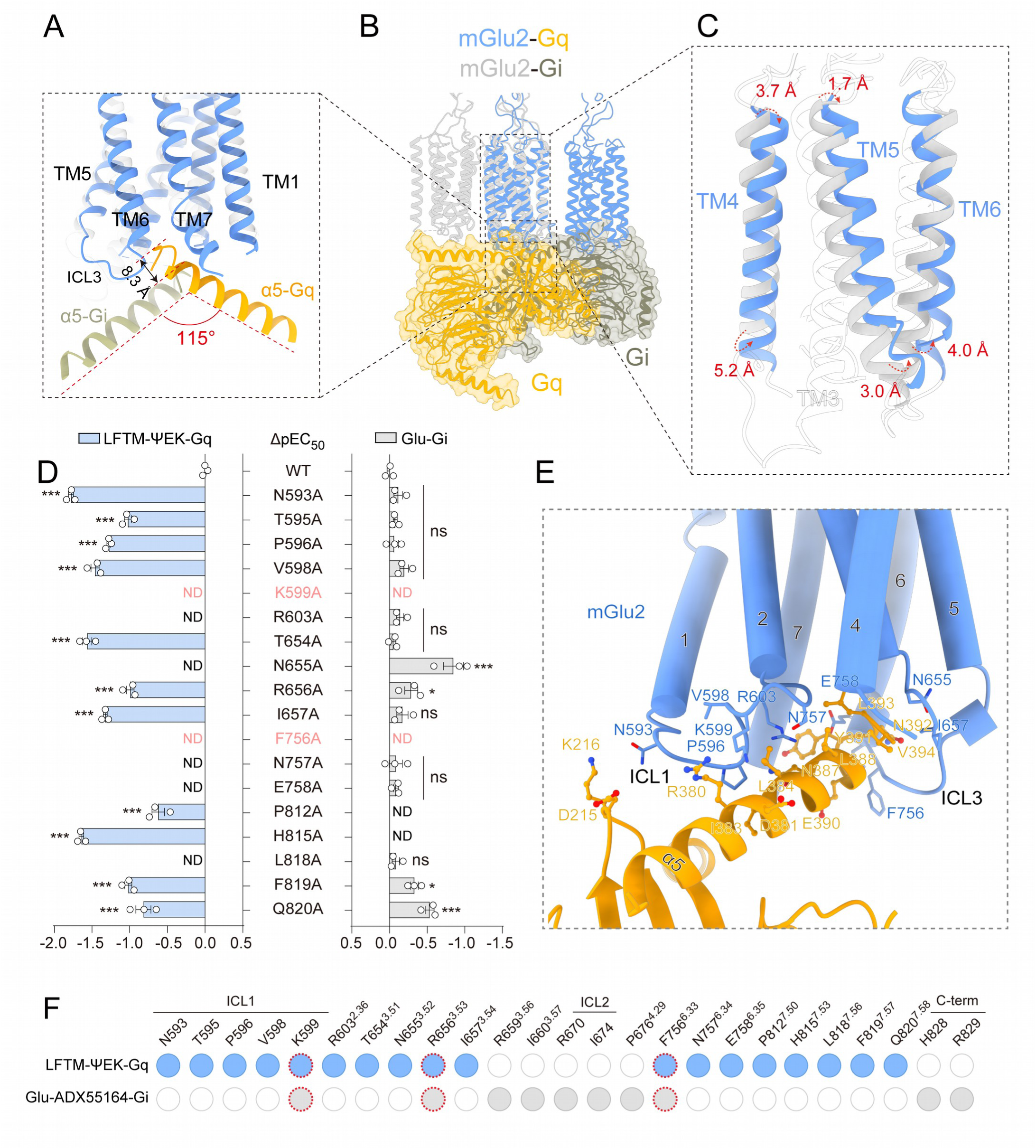
Distinct Gq coupling mode of mGlu2. **(A-C)** The LFTM-ΨEK–mGlu2–Gq and Glu–ADX55164–mGlu2–Gi (PDB ID: 7MTR) complexes were aligned along TM2 and TM3 of G protein-coupled protomers. (LFTM-ΨEK-mGlu2 dimer, blue; Glu-ADX55164-mGlu2, gray; Gq protein, Amber Glow; Gi protein, brown). (A) Detailed representation of the receptor-binding orientation of the α5-helices from Gq versus Gi proteins in mGlu2-Gq and mGlu2-Gi complexes. (B) Overall view of the structural alignment between the LFTM-ΨEK–mGlu2–Gq and Glu–ADX55164–mGlu2–Gi complexes. (C) Detailed representation of transmembrane helix conformational differences in G protein-bound protomers between mGlu2-Gq and mGlu2-Gi complexes. **(D)** Effects of different mutations within the G protein interface of LFTM-ΨEK-mGlu2-Gq complex on LFTM-ΨEK-induced Gαq-Gγ dissociation and Glu-induced Gαi-Gγ dissociation in mGlu2 overexpressing cells. **(E)** Schematic diagram of the interface between Gq and mGlu2, involving ICL1, TM3, TM6, and TM7. LFTM-ΨEK-mGlu2 dimer, blue; Gq protein, Amber Glow. **(F)** Comparison of residues in the binding pockets for G protein in the LFTM-ΨEK-Gq and Glu-ADX55164-Gi complexes. The residues that engage with Gq and Gi are shown in blue and gray, respectively. The residues, which interacts with both Gq and Gi, are shown in red dashed borders. (D) Values are shown as the mean ± SEM of three independent experiments (n=3). \**P* < 0.05; \*\*\**P* < 0.001; ns, no significant difference; ND, not detected, the mGlu2 mutation groups compared with the WT groups. All the data were analyzed by one-way ANOVA with Dunnett’s multiple comparisons test.

Among 20 residues composing the Gq interface of mGlu2, only 3 are identical with those in the glutamate-ADX55164-mGlu2-Gi interface, suggesting distinct G protein coupling mechanisms (Figure 7F). In the former complex, the residues T595^ICL1^-P596 ^ICL1^-V598^ICL1^ form extensive hydrophobic interactions with α5 helix of Gq (Figure 7E). N593 ^ICL1^ and R603^2.36^ establish hydrogen bond networks with D215^G.s2s3.01^-K216^G.s2s3.02^ and N387^G.H5.19^, respectively, whereas K599^ICL1^ forms a charge interaction with D381^G.H5.13^ (Figure 7E). Additionally, the hydrophobic cavity formed by TM3/6/7 further envelops the helical terminus of α5 helix of Gq (Figure 7E). Specifically, residues I657^3.54^-N655^3.52^-T654^3.51^ in TM3 establish a hydrophobic interaction network with L393^G.H5.25^ -V394^G.H5.26^ of the α5-helix of Gq (Figure 7E). In TM6, E758^6.35^ and N757^6.34^ form hydrogen bonds with N392^G.H5.24^ and E390^G.H5.22^, respectively (Figure 7E). Meanwhile, the aromatic triad Y391^G.H5.23^ and F819^7.57^ in TM7 stabilizes the mGlu2-Gq complex via π-π stacking interactions. Mutations of residues along these Gq interface, such as N593^ICL1^A, V598^ICL1^A, R603^2.36^A, T654^3.51^A, N655^3.52^A and E758^6.35^A, significantly impair the force-stimulated mGlu2-Gq signal, but not Gi signal in response to glutamate stimulation (Figure 7D-E and Extended Data Figure 10F-H). Collectively, these biochemical data supported the structural observation of distinct coupling mechanisms between mGlu2 and Gq or Gi.

### Propagating pathway underlies force-induced mGlu2-Gq coupling

Comparison of the structures of LFTM-ΨEK-mGlu2-Gq and mGlu2^apo^ allows us to propose a possible mechanism for mGlu2 activation in response to LFTM-ΨEK binding (Figure 8C). The engagement of Y787^7.25^ with the E558^LFTM-ΨEK^ of LFTM-ΨEK via a hydrogen bond and the interaction between R788^7.26^ and T792^7.30^ and D565-R562 of LFTM-ΨEK caused the side chain of Y787^7.25^ to flip counterclockwise by approximately 60° and the side chain of R788^7.26^ rotates clockwise by approximately 30° (Figure 8B). Accompanied with these changes, T792^7.30^, C795^7.33^, and V796^7.34^ shift towards TM1 by approximately 2.2∼3.4 Å, resulting in a 3.5 Å upward shift of TM7 (Figure 3E and 8A-B). These conformational rearrangements cause an 86° rotation of M794^7.32^ that disrupts the native M794^7.32^-F780^6.57^-F776^6.53^ hydrophobic triad core in mGlu2^apo^ structure (Figure 8D). Disruption of this hydrophobic network drives a roughly 102° rotation of the toggle switch element W773^6.50^, enabling its indole ring to form a hydrogen bond with S801^7.39^ and to stabilize the TM6 orientation in the active state (Figure 8D). The S801^7.39^ then builds up a hydrogen bond with the T769^6.46^, and connects to intracellular ends of TM6-TM7 residues via a conformational propagating path of V804^7.42^-T765^6.42^-C808^7.46^-I762^6.39^-F761^6.38^ (Figure 8E). Structural alterations of these path residues in response to LFTM-ΨEK binding caused a displacement of approximately 3 Å in the intracellular end of TM6 towards TM7, and the side chains of E758^6.35^, F756^6.33^, N757^6.34^, H815^7.53^ to rotate toward the intracellular cavity of the 7TM bundle, enabling Gq engagement (Figure 8F). By forming W773^6.50^-S801^7.39^-T769^6.46^ polar interaction network, the conformational change of extracellular end of TM7 is transmitted to TM6, causing a displacement of approximately 3.7 Å in TM6 towards TM7 (Figure 3E and 8D-E). Alanine mutation scanning data supported the importance of these propagation path residues in mediating LFTM-ΨEK induced mGlu2 activation (Figure 8G and Extended Data Figure 11A-D).

**Figure 8.**
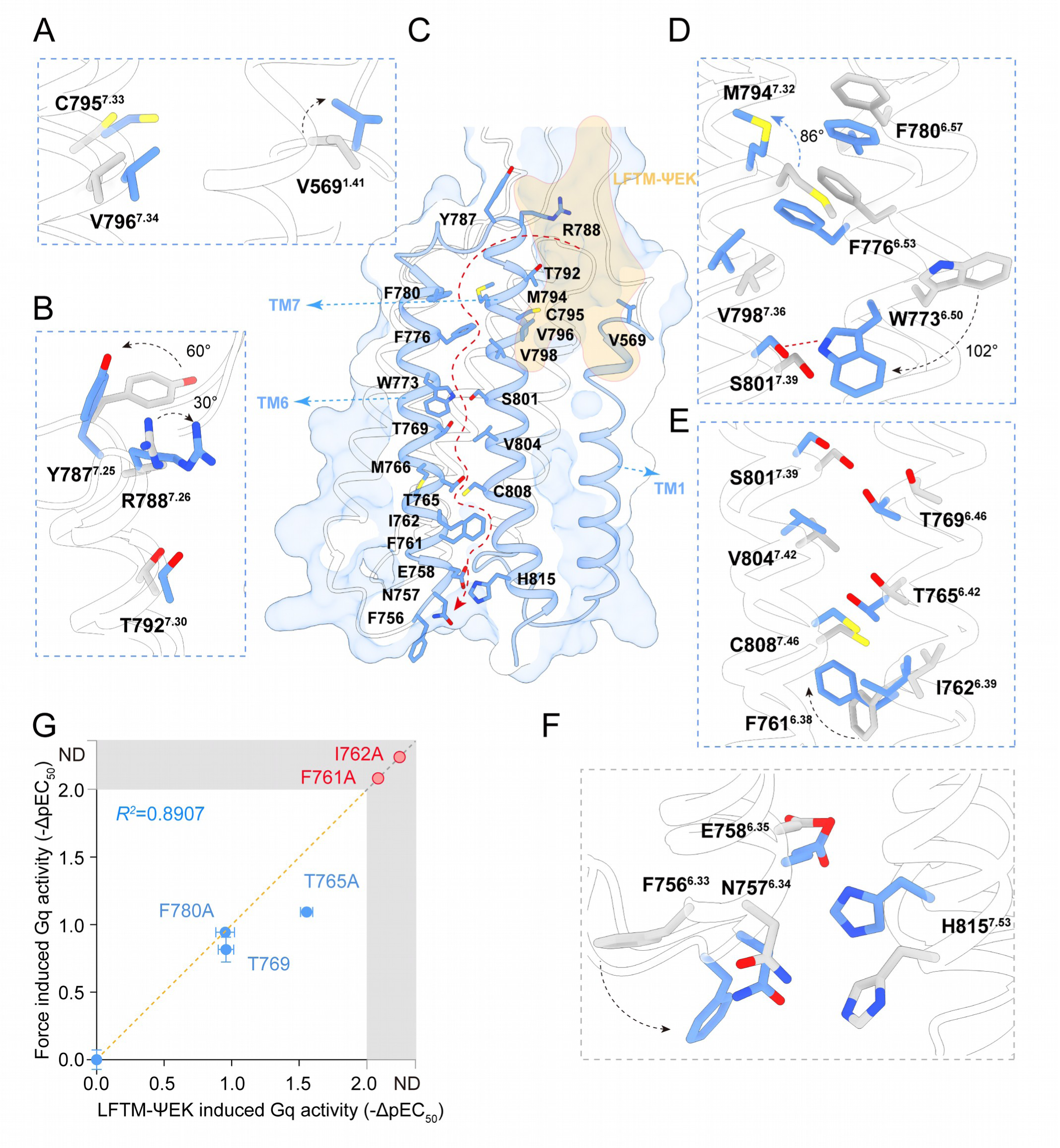
Propagating path underlining force-induced mGlu2 coupling to Gq. **(A-F)** Schematic illustration propagating path of mGlu2 receptor activation in response to LFTM-ΨEK peptide (C). Compared with the mGlu2 apo state (gray), the active state of LFTM-ΨEK-mGlu2-Gq complex (blue) exhibits conformational shifts of key residues in the ligand-binding pocket (V569^1.41^, Y787^7.25^, R788^7.26^) (A-B), which induce rearrangement of the hydrophobic interaction network involving F780^6.57^-M794^7.32^-F776^6.53^ (D). This leads to the rotation of the toggle switch residue W773^6.50^ and subsequent formation of a hydrogen-bond interaction with S801^7.39^, ultimately driving conformational changes in the intracellular ends of TM6 and TM7 helices to facilitate Gq protein coupling (E-F). **(G)** Summary of the effects of mutations of propagating path residues on the LFTM-ΨEK- and force-induced Gq activity measured by BRET assay. The X axis represents -ΔpEC_50_ values between LFTM-ΨEK-induced Gq activity of mGlu2 mutants and mGlu2-WT; positive values indicate reduced potency of LFTM-ΨEK. The Y-axis represents -ΔpEC_50_ values between force-induced Gq activity of mGlu2 mutants and mGlu2-WT; positive values indicate reduced potency of force. Mutants in the first quadrant displayed reduced LFTM-ΨEK-induced Gq activity and diminished force-sensing potential, mimicking the coordinated changes in LFTM-ΨEK- and force-induced mGlu2 Gq activity modulation (n=3). The orange dashed line represents the y=x reference, while the R² value reflects the correlation between mutant-induced differences in LFTM-ΨEK- and force-stimulated Gq activity changes.

### Structural integrity of mGlu2 mechanosensitivity in kinocilium, but not glutamate binding, is required for normal vestibular functions

To further validate the mechanism underlying the mechanosensitivity of mGlu2 and to explore its physiological significance in equilibrioception, we performed an *in vivo* rescue experiment. We chose the mGlu2^Δ1-555^ construct that presented mechanosensitivity comparable to that of WT mGlu2 but completely abolished the response to glutamate stimulation due to the lack of N-terminal glutamate-binding domains. We further generated a mutant, V598A, which presented cell surface expression similar to that of mGlu2^Δ1-555^ but nearly completely abolished the mechanosensitivity (Figures 9A and 9B). We packaged the coding sequences of mGlu2^Δ1-555^ or the force-insensitive mutant into a modified AAV-ie-*Grm2pr*-EGFP vector, which enabled the protein expression driven by endogenous mGlu2 promotor. The virus was delivered into the utricular hair cells of *Pou4f3-CreER^+/−^; mGlu2^fl/fl^* mice, in which the mGlu2 was specifically ablated in inner ear hair cells, at P3 through round window membrane injection (Figure 9C). The mGlu2 was found to be re-expressed at the kinocilium of the utricular hair cells of the *mGlu2*-deficient mice 5 weeks after the administration of AAV-ie encoding mGlu2^Δ1-555^ or the mGlu2^Δ1-555-V598A^ mutant, as shown by immunofluorescence analysis (Figures 9D and 9E). As a negative control, the *Pou4f3-CreER^+/−^; mGlu2^fl/fl^*mice infected with AAV-ie-*mGlu2*-EGFP exhibited no detectable mGlu2 expression in kinocilium (Figure 9F).

**Figure 9.**
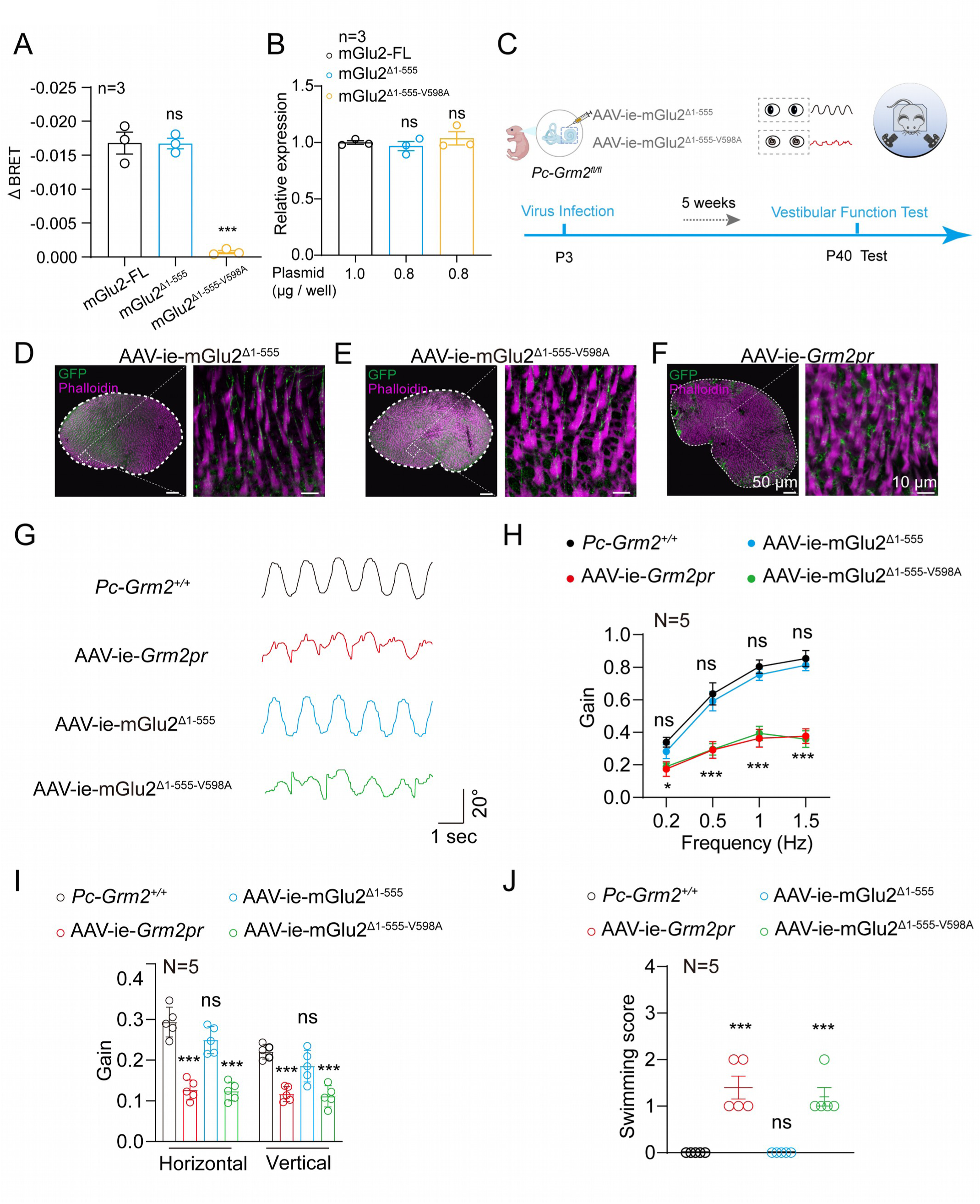
Force-sensitive mGlu2 mutant expression in hair cells of *Grm2*-deficient mice prevents vestibular dysfunction. **(A)** Force-induced Gq activation in HEK293 cells transfected with full-length mGlu2 or its N-terminal truncated mutants, including mGlu2^Δ1-555^, mGlu2^Δ1-555-V598A^ (n=3). **(B)** Equal cell surface expression levels of full-length mGlu2, mGlu2^Δ1-555^ and mGlu2^Δ1-555-V598A^ in HEK293 cells measured by cell-surface ELISA assay (n = 3). **(C)** Schematic representation of the strategy to reintroduced mGlu2 or its mutants expression in the utricular hair cells of the *Pc-Grm2^fl/fl^* mice via AAV-ie delivery and the vestibular function validation. **(D-F)** Expression of mGlu2-GFP (green) with the stereocilia (magenta) in utricular hair cells derived from AAV-ie-mGlu2^Δ1-555^-treated *Pc-Grm2^fl/fl^* mice (referred to as AAV-ie-mGlu2^Δ1-555^ mice) (D), AAV-ie-mGlu2^Δ1-555-V598A^-treated *Pc-Grm2^fl/fl^* mice (referred to as AAV-ie-mGlu2^Δ1-555-V598A^ mice) (E) and AAV-ie-*Grm2pr*-treated *Pc-Grm2^fl/fl^* mice (referred to as AAV-ie-*Grm2pr* mice) (F). Scale bars: 50 μm and 10 μm for low and high magnification view, respectively. **(G-H)** Representative recording curves (**G**) and quantification of the VOR gain values (**H**) of *Pc-Grm2^+/+^* mice, AAV-ie-mGlu2^Δ1-555^ mice, AAV-ie-mGlu2^Δ1-555-V598A^ mice and AAV-ie-*Grm2pr* mice in response to earth-vertical axis rotation (N=5 mice per group). **(I)** Quantification of the VOR gain values of *Pc-Grm2^+/+^* mice, AAV-ie-mGlu2^Δ1-555^ mice, AAV-ie-mGlu2^Δ1-555-V598A^ mice and AAV-ie-*Grm2pr* mice in response to off-vertical axis rotation (N = 5 mice per group). **(J)** Quantification of the swimming scores of *Pc-Grm2^+/+^* mice, AAV-ie-mGlu2^Δ1-555^ mice, AAV-ie-mGlu2^Δ1-555-V598A^ mice and AAV-ie-*Grm2pr* mice (N = 5 mice per group). (A-B) ***P<0.001; ns, no significant difference. N-terminal truncated mutants of mGlu2 compared with full-length mGlu2. (H-J) *P < 0.05; ***P < 0.001; ns, no significant difference. AAV-ie-mGlu2^Δ1-555^ mice, AAV-ie-mGlu2^Δ1-555-V598A^ mice or AAV-ie-*Grm2pr* mice compared with *Pc-Grm2^+/+^* mice. The bars indicate mean ± SEM values. All data were statistically analyzed using one-way (A, B, I, J) or two-way (H) ANOVA with Dunnett’s post hoc test.

We next investigated the vestibular function of the AAV-ie-treated mice by examining their vestibular-ocular reflex (VOR) during sinusoidal head rotations. Notably, the *mGlu2*-deficient mice treated with AAV-ie encoding mGlu2^Δ1-555^ showed significantly elevated VOR gain values at all tested frequencies in both earth-vertical and off-vertical axis rotation tests, which were restored to levels comparable to those of the control *Pou4f3-CreER^+/−^; mGlu2^+/+^* mice (Figures 9G-9I). Consistently, the AAV-ie-mGlu2^Δ1-555^-treated mice exhibited improved performance in the forced swimming test, which was also comparable to that of the *Pou4f3-CreER^+/−^; mGlu2^+/+^* mice (Figure 9J). In contrast, the *mGlu2*-deficient mice administered with AAV-ie encoding force-insensitive mGlu2^Δ1-555-V598A^ mutant behaved similarly as the negative control mice treated with AAV-ie-*mGlu2pr*-EGFP, showing no significant improvement in any of the above vestibular functional tests (Figures 9G-9J). These findings suggest that the mechanosensitivity of mGlu2 expressed at the kinocilium of vestibular hair cells is required for its regulation of normal equilibrioception.

## Discussion

In our parallel study and current work, we have identified that a Class C GPCR, mGlu2, is able to sense force and transduce mechanical stimulation to Gq coupling. Importantly, this physical signal induced signaling bias is distinct from the well-characterized Gi signaling stimulated by endogenous agonist of mGlu2, the glutamate. Whereas mGlu2 activated by glutamate and its Gi coupling play important roles in neurotransmitter release ^37^, the forced-induced mGlu2-Gq coupling is essential for normal balance. These observations suggested that diverse ligand recognition in particular different chemical (glutamate) or physical (Force) kinds and distinct G protein subtype signaling underlies mGlu2 functions in different physiological contexts. In the classic glutamate-mGlu2-Gi signaling, the glutamate acts as an orthosteric ligand binding within the VFT domains of both protomers while the allosteric ligands bind randomly to the TMD region (between TM5-6-7) of either protomer. In contrast, we found that mechanical force, a kind of physical stimulation, acting as a compressive signal, activates mGlu2 through a specific FTM motif at the N-terminus of the 7TM domain in a single mGlu2 protomer, independent of the VFT domains. This mechanism enables us to recall the force-induced aGPCR activation, which is dependent on the “*Stachel* peptide”^15, 16^. Moreover, the FTM sequence binds within a shallow pocket between TM1 and TM7, which is distinct from the traditional allosteric PAM binding site^25, 29^.

Compared with apo-mGlu2 structure, glutamate binding induces VFT domain closure and rotation, driving the 7TMs from an inactive conformation characterized by a TM3/4 interface symmetry to an active, asymmetric conformation featured with a TM6/6 interface between two protomers. Allosteric agonists further stabilize this asymmetric state. Conversely, during force activation of mGlu2-Gq signaling, the VFT domains on both protomers remain largely open without significant rotation, indicating their dispensable roles in mechanotransduction. However, we observed that VFT domains might undergo a small-angle deflection induced by FTM motif binding, driving the protomers apart and forming an asymmetric interface characterized by TM3-4/TM4. This unique micro-conformational change pattern may underlie the ultra-high sensitivity of the vestibular system (Zhou et al., 2025, paralleling submitted study). Our structural observation, conformational change analysis by BRET and mutational analysis further suggest that the mechanical signal is transmitted through the FTM motif to the 7TM of a single protomer of mGlu2. This is significantly different from the activation mode in the mGlu2-Gi pathway, where the signal is transmitted to the TMD through a concerted twist of both extracellular domains. The FTM motif binds within a shallow pocket formed by TM1-TM7-ECL2. Through a hydrophobic interaction network involving M794^7.32^-F780^6.57^-F776^6.53^, termed the hydrophobic triad motif (HTM), the peptide binding or force signal is relayed to the “toggle switch” residue W773^6.50^, which then activates downstream pathways via an S801^7.39^-T769^6.46^ hydrogen bond network and a V804^7.42^-T765^6.42^-C808^7.46^-I762^6.39^-F761^6.38^ conformational propagation pathway. This force transmission mechanism contrasts with the traditional mode where chemical agonists induced activation of class C GPCRs^25–29, 36, 38–46^.

Notably, within the FTM-mGlu2-Gq complex, the Gq protein specifically binds to the ICL1-TM3-TM6-TM7 pocket of the PB protomer. Its α5 helix couples in a unique conformation, deeply inserted and oriented towards TM6—spatially opposing the TM3-ICL2-ICL3 binding mode of the Gi α5 helix in Glu-mGlu2-Gi complex structure, forming a near antiparallel arrangement. Gq engages deeper into the transmembrane bundle (8.3 Å shift) compared to Gi, forming a new interface dominated by ICL1 interactions (T595^ICL1^-P596^ICL1^-V598^ICL1^). Only 15% of residues overlap between Gq and Gi binding sites, highlighting distinct coupling mechanisms. Examination of mutation effects further support that the Gq and Gi coupled to mGlu2 through distinct modes (Figure 10).

**Figure 10.**
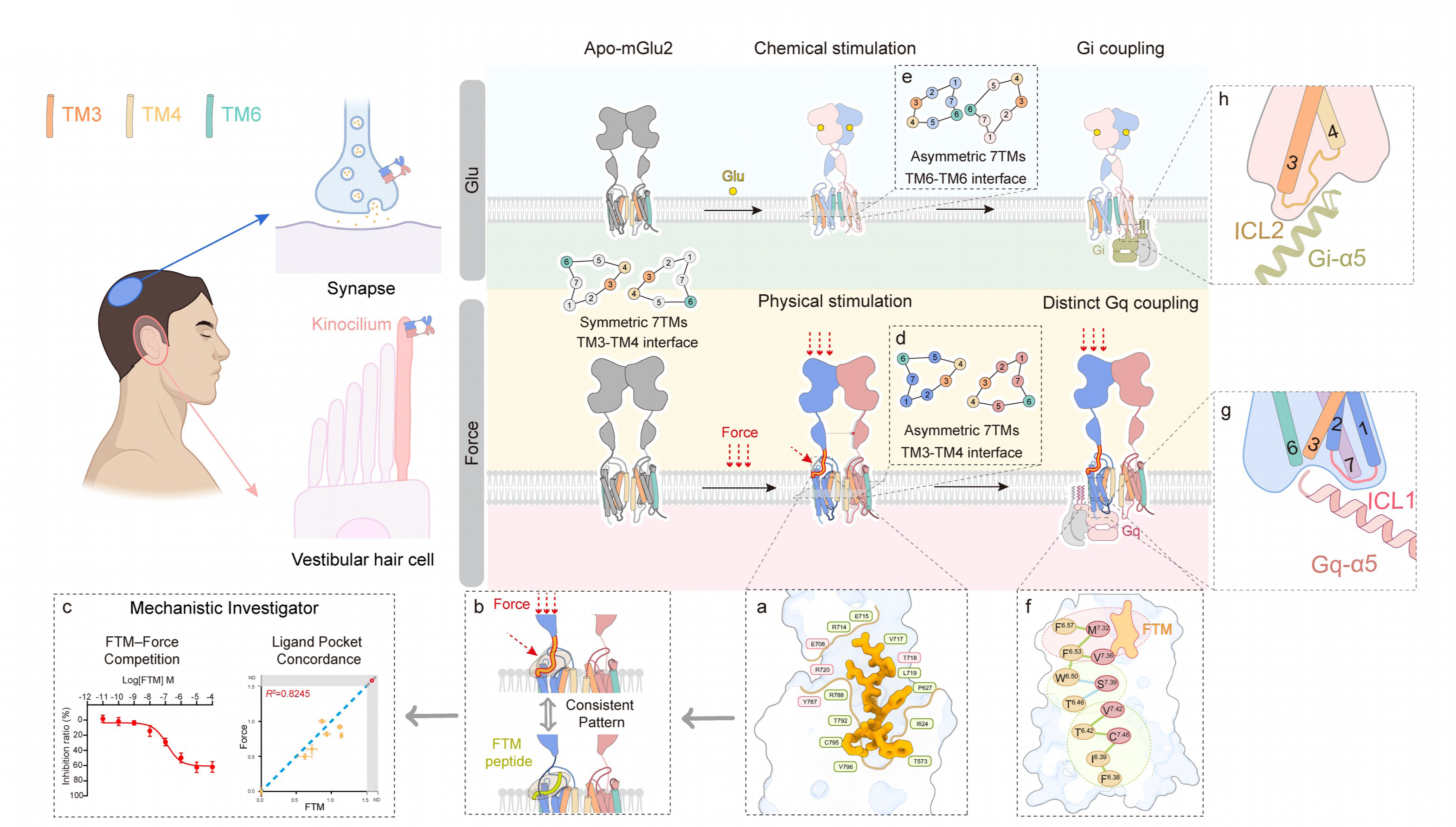
Models describing the distinct activation modes of mGlu2 in response to chemical (glutamate) and physical (force) stimulation in different physiological contexts. In its apo state, mGlu2 adopts a symmetric conformation characterized by open VFTs, distant CRDs, and a symmetric 7TM dimer with a TM3–TM4 interface. The mechanical force induces mGlu2 activation by propelling the FTM motif into an allosteric pocket composed by TM1, TM7, and ECL2 (a). FTM-derived peptide is able to mimic the force-induced mGlu2 activation by binding to the same pocket (b). This conclusion is supported by the following evidences: (1) the FTM peptide competitively antagonizes the force-induced mGlu2 activation, and (2) the mutational effects on pocket residues are similar between force- and FTM-induced mGlu2 activation (c). Upon force stimulation, the 7TMs of mGlu2 undergoes a slight angular shift, forming an asymmetric TM3–TM4 interface (d). In contrast, the glutamate binds to the VFT of mGlu2, driving the 7TMs to from an active, asymmetric conformation featured with a TM6-TM6 interface (e). The mechanical signal is transmitted from the FTM to the 7TM of a single protomer of mGlu2, through a hydrophobic triad motif and a TM6–7 conformational pathway (f). This leads to a specific Gq coupling to mGlu2 via an ICL1-TM3-TM6-TM7 interface, allowing a deeper insertion of the α5 helix (∼8.3 Å) and nearly a 180° turn in the insertion direction of the α5 helix compared to that of Gi coupling which selectively engages ICL2-TM3 domains of mGlu2 (g-h).

Evolutionary analysis of the FTM motif provides a molecular explanation for the uniqueness of force induced activation and Gq coupling of mGlu2, but not other 7 mGlu members. The core residues of the FTM motif, P^557^XE^559^YXRW^563^, are highly conserved across vertebrates but not in other mGlu family members. This explains the exclusive mechanosensitivity of mGlu2 acquired during species evolution. Furthermore, we revealed that synthetic FTM peptides might mimic mechanical force to activate Gq and competitively inhibit the response to force stimulation (EDF Fig. 2I), confirming FTM as the core element for mechanical force transduction. Through unnatural amino acid substitution and sequence optimization based on species variants, we screened for a higher-affinity peptide agonist, LFTM-ΨEK. In addition to synthetic compounds, these discoveries suggested peptide-based agonists could be developed to modulate mGlu activities. Finally, using *Pou4f3-CreER^+/−^; mGlu2^fl/fl^* mice supplemented with AAV-ie encoding mGlu2^Δ1-555^ or the mGlu2^Δ1-555-V598A^ mutants, we were able to provide in vivo evidence that structural integrity of force sensing, but not glutamate sensing, is required for normal balance.

Collectively, we discovered that force and peptide can activate mGlu2, a class C GPCR only known to sense neurotransmitter in previous studies, and revealed unique structural mechanisms underlying force- or peptide-induced mGlu2 activation, G protein subtype coupling, and propagating path underlying conformational changes

## Acknowledgments

This work was supported by the National Key R&D Program of China (2023YFA1801100 and 2024YFA1107500 to Z.Y.; 2024YFA1802900 to J.C.), the Foundation for Innovative Research Groups of the National Natural Science Foundation of China (T2321004 to J.-P.S.), the National Science Fund for Distinguished Young Scholars grant (82425105 to J.-P.S. and 82225011 to X.Y.), the National Science Fund for Excellent Young Scholars (82422072 to Z.Y.), the National Natural Science Foundation of China (32130055, 32361163612, and 82330118 to J.-P.S.; 92357303 to X.Y.; 824B2018 to W.-F.Z.), the International (Regional) Cooperation and Exchange Programs of the National Natural Science Foundation of China (3231101039 to J.-P.S.), the Taishan Scholars Program (tsqn202408028 to Z.Y.), the Key Research and Development Program of Shandong Province (2021CXGC011105 to J.-P.S.), and the Unconventional Innovation Project of Beijing Natural Science Foundation (F251012 to J.-L.W.). J.-P.S. is supported by the Tencent New Cornerstone Investigator Program and Open Research Project in State Key Laboratory of Vascular Homeostasis and Remodeling (Peking University). We would like to thank the Peking University Health Science Center Cryo-Electron Microscopy Facility for our apo-mGlu2 dimer, FTM-mGlu2 dimer, LFTM-ΨEK-mGlu2 dimer, and LFTM-ΨEK-mGlu2-Gαq complex.

## Author contributions

J.-P.S. conceptualized, organized, and supervised the entire study. J.-P.S. designed the research strategy for the force-sensing mechanism of mGlu2, as well as the pharmacology, cell signaling, and structural studies. J.-F.L. conceived the method to validate the mGlu2 dimer interaction interface and designed cross-linking and fluorescent-labeled blot experiments. Z.Y. and X.Y. conceived the animal behavioral method to validate mice balance perception abilities and designed functional assays. J.-P.S., Z.Y., J.-F.L., and X.Y. participated in data analysis and interpretation. J.-H.D. and W.-P.Z. established the mGlu2 293F cell expression construct, developed purification protocols for the apo-mGlu2 dimer, FTM-mGlu2 dimer, LFTM-ΨEK-mGlu2 dimer, and LFTM-ΨEK-mGlu2-Gαq complex, and prepared cryo-EM samples. J.-L.W., J.-H.D., and W.-P.Z. prepared cryo-EM grids. W.D., J.-L.W., and J.-H.D. performed cryo-EM map calculation, model building, and structural refinement. S.-H.Z. and Q.-Y.Z. conducted animal experiments and performed related data analysis. W.-F.Z., W.-P.Z. and J.C. performed BRET dissociation assays and analyzed the relevant data. W.-F.Z. and J.-H.D. performed flash-BRET signal detection and analyzed the relevant data. C.-J.X. performed crosslinking and fluorescently tagged blotting experiments and analyzed the relevant data. C.Z. and Y.-N.S. performed molecular dynamics simulation predictions. J.-P.S., Z.Y., J.-H.D., J.-L.W., and S.-H.Z. wrote the manuscript. J.-F.L., X.Y., W.-F.Z., W.-P.Z., R.-J.C. and Y.S. provided important discussions and essential revisions. J.-H.D. and W.-F.Z. provided Figures 1, 2, 3, 4, 5, 6, 7, and 8. C.-J.X. provided Figure 4. S.-H.Z. and Q.-Y.Z. provided Figure 9. J.-H.D., W.-F.Z., W.-P.Z., and C.-J.X. provided extended data figures.

